# Allicin inhibits SARS-CoV-2 replication and abrogates the antiviral host response in the Calu-3 proteome

**DOI:** 10.1101/2021.05.15.444275

**Authors:** Kirstin Mösbauer, Verena Nadin Fritsch, Lorenz Adrian, Jörg Bernhardt, Martin Clemens Horst Gruhlke, Alan John Slusarenko, Daniela Niemeyer, Haike Antelmann

## Abstract

The Severe Acute Respiratory Syndrome Coronavirus 2 (SARS-CoV-2) pandemic is a major health burden. Volatile garlic organosulfur compounds, such as the thiol-reactive allicin (diallyl thiosulfinate) exert strong antimicrobial activity against various respiratory pathogens. Here, we investigated the antiviral activity of allicin against SARS-CoV-2 in infected Vero E6 and Calu-3 lung cells. Calu-3 cells showed greater allicin tolerance due >4-fold increased GSH levels compared to Vero E6. However, biocompatible allicin doses efficiently inhibited viral replication in both cell lines. Proteome analyses of SARS-CoV-2 infected Calu-3 cells revealed a strong induction of the antiviral interferon-stimulated gene (ISG) signature (e.g. cGAS, Mx1, IFIT, IFIH, IFI16, IFI44, 2’5’OAS and ISG15), pathways of vesicular transport, tight junctions (KIF5A/B/C, OSBPL2, CLTC1, ARHGAP17) and ubiquitin modification (UBE2L3/5), as well as reprogramming of host metabolism, transcription and translation. Allicin abrogated the ISG host response and reverted the host cellular pathways to levels of uninfected Calu-3 cells, confirming the antiviral and immunomodulatory activity of allicin in the host proteome. Thus, biocompatible doses of allicin could be promising for protection of lung cells against SARS-CoV-2.

## Introduction

The Severe Acute Syndrome Coronavirus 2 (SARS-CoV-2) causes Coronavirus disease (COVID-19), which represents a global health burden (Zhou et al 2020). COVID-19 is often associated with immunopathology since severely ill patients had decreased levels of T lymphocytes, including regulatory T cells, cytotoxic and helper T cells and natural killer cells (Donma & Donma 2020, Qin et al 2020, Wei et al 2020). Patients with severe illness showed a cytokine storm syndrome associated with a dysregulated immune activation and hyperinflammation (Fara et al 2020). High levels of pro-inflammatory cytokines IL-1ß, IL-2, IL-6, IL-7, IL-10, macrophage inflammatory protein-1A (MIP-1A), TNF-α and INF-γ have been detected, connecting the uncontrolled inflammation and dysregulation of the immune response with the high mortality in severely ill COVID-19 patients (Fara et al 2020, Qin et al 2020, Wei et al 2020). While mild infections were characterized by highly activated HLA-DR^hi^CD11c^hi^ inflammatory monocytes with the interferon-stimulated gene (ISG) signature, severe illness was manifested by dysfunctional neutrophil precursors, and HLA-DR^lo^ monocytes with pro-inflammatory functions (Schulte-Schrepping et al 2020). These immunological markers of pro-inflammatory cytokines and the dysfunctional myeloid compartment might help to identify drug targets to prevent progression to severe illness (Fara et al 2020, Schulte-Schrepping et al 2020).

While global vaccination campaigns are underway, the development of efficient therapies to prevent COVID-19 disease progression is an urgent need. FDA-approved repurposed COVID-19 drug candidates include for example the antivirals remdesivir, lopinavir/ritonavir and favipiravir, which are used for the treatment of other viral infections, such as hepatitis, HIV and influenza, respectively (Drozdzal et al 2020). Further clinical trials are investigating the use of antibodies against IL-6 (tocilizumab) and CD-147 (meplazumab) or chloroquine and hydroxychloroquine, which inhibit T-cells and decrease pro-inflammatory cytokines (Drozdzal et al 2020).

Apart from drugs, functional food based on herbal medicine might be supportive due to their immunomodulatory, antioxidant, anticancer, antimicrobial and antiviral activities (Donma & Donma 2020, Khubber et al 2020). For example, garlic plants (*Allium sativum*) produce volatile organosulfur compounds, such as diallyl thiosulfinate (allicin) and diallyl polysulfanes, which are known to stimulate the immune system by modulation of cytokine secretion and pro-inflammatory cytokines (Borlinghaus et al 2014, Donma & Donma 2020, Khubber et al 2020). Allicin showed broad-spectrum antimicrobial activity against several pathogenic bacteria, viruses, fungi and parasites and has been used for the treatment of pneumonia, tuberculosis and the common cold since ancient times (Arbach et al 2019, Block 2010, Borlinghaus et al 2014, Münchberg et al 2007, Rabinkov et al 1998, Reiter et al 2017, Rivlin 2001; 2006, Rouf et al 2020). Administration of the volatile allicin by inhalation can efficiently reach the infected lung tissue. Already in 1927, vapor from garlic extracts was supplied in face masks by Minchin’s inhaler and used to treat pulmonary tuberculosis (Minchin 1927, Reiter et al 2017).

Allicin is a strongly thiol-reactive compound, which reacts with Cys thiols via thiol-disulfide exchange reactions, leading to *S*-thioallylations of proteins (Miron et al 2010, Miron et al 2000). Widespread *S*-thioallylations of redox-sensitive Cys residues in proteins were identified in the proteome of human Jurkat cells, *E. coli*, *S. aureus* and *B. subtilis* (Chi et al 2019, Gruhlke et al 2019, Loi et al 2019, Miron et al 2010, Müller et al 2016, Rabinkov et al 1998). In Jurkat cancer cells, 332 *S*-thioallylated proteins were identified 10 min after allicin treatment, including highly abundant cytoskeleton proteins, HSP90 chaperones, translation elongation factors and glycolytic enzymes. Allicin caused disruption of the actin cytoskeleton, enzymatic inactivation and Zn^2+^ release to stimulate the IL-1-dependent IL-2 secretion by T-cells as an immunomodulatory effect (Gruhlke et al 2019).

In addition, *S*-thioallylations deplete low molecular weight thiols, such as glutathione (GSH) and bacillithiol (BSH) in bacteria and yeast cells (Arbach et al 2019, Gruhlke et al 2010, Gruhlke et al 2019). Thus, allicin leads to oxidative stress responses, inhibition of protein functions and an impaired cellular redox balance. Since SARS-CoV-2 is rich in Cys residues in its surface spike glycoprotein, a reduced state of the host cell cytoplasm is required for efficient virus entry and membrane fusion. Moreover, allicin is cell permeable and has been shown to cause transient pore formation in phospholipid membranes, which may contribute to the killing of SARS-CoV-2 by affecting its envelope (Gruhlke et al 2015, Miron et al 2000). While the antiviral effect of allicin has been studied against several viruses that cause respiratory tract infections, including influenza, SARS-CoV and rhinovirus (Rouf et al 2020), mechanistic insights on its proposed antiviral effects against SARS-CoV-2 in the infected host cell are lacking.

In this work, we show that allicin at biocompatible doses efficiently inhibits replication of SARS-CoV-2 in the primate kidney-derived cell line Vero E6 and the human lung cell line Calu-3. We further identified proteome changes caused by SARS-CoV-2 infection and the effect of allicin on these host pathways. While the interferon-stimulated gene (ISG) signature was most prominently upregulated in SARS-CoV-2 infected Calu-3 cells, the ISG response and other host cellular pathways were reverted to Mock levels by allicin. Thus, allicin exerts a beneficial effect as an antiviral and immunomodulatory compound in cell lines and could be utilized as a supportive therapy for the treatment of COVID-19.

## Results

### Biocompatible allicin concentrations in Vero E6 and Calu-3 cells correspond to the intracellular GSH levels

High allicin doses were previously shown to act as irritant and cause cellular damage (Bautista et al 2005, Hitl et al 2021). Thus, a cell viability assay was used to determine the biocompatible, non-harmful doses of allicin in Calu-3 and Vero E6 cells. Both cell lines differed strongly in their susceptibilities towards allicin. Calu-3 cells showed high viability rates of ~ 85% after treatment with 200 μM allicin. Even concentrations of 300 μM allicin decreased the viability rate of Calu-3 cells only non-significantly to ~70% **(Fig. 1A)**. Treatment of Vero E6 cells with 75 μM allicin led to a cell viability rate of 86% **(Fig. 1B)**, whereas 150 μM allicin resulted in killing of 99 % of the cells. Thus, the sub-lethal biocompatible doses of allicin were determined as 50-75 μM in Vero E6 cells and 100-200 μM in the more tolerant Calu-3 cells.

**Fig. 1.**
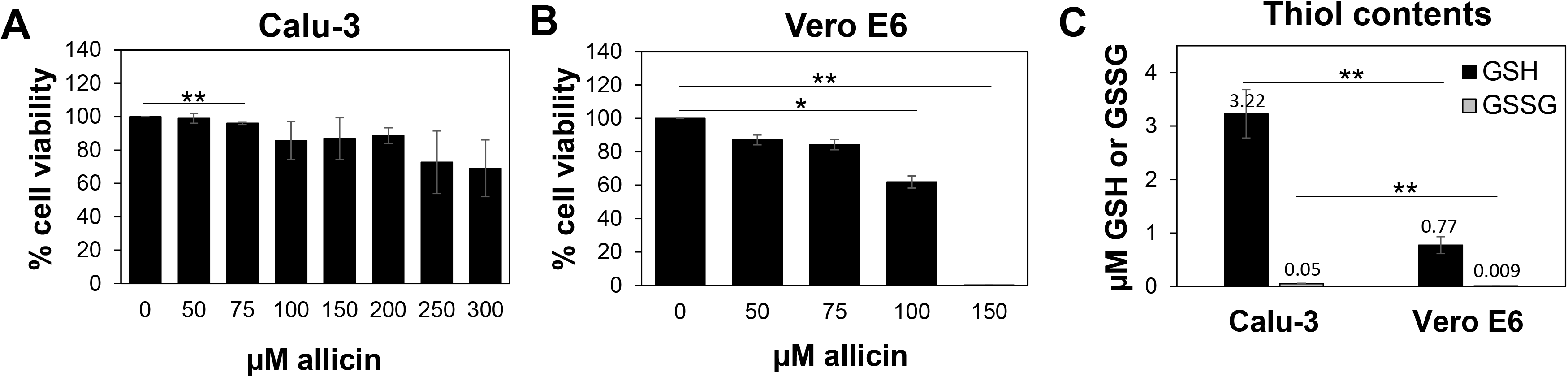
Human Calu-3 cells are more resistant to allicin compared to Vero E6. **(A,B)** Cell viability of untreated and allicin treated Calu-3 **(A)** and Vero E6 cells **(B)** were measured after 24 h using the CellTiter-Glo Luminescent Cell Viability Assay (Promega) according to the manufacturer’s instructions. Calu-3 cells were not affected by exposure to ≤ 200 μM allicin, while concentrations of ≥100 μM allicin interfered with Vero E6 cell viability. The viability of the control without allicin was set to 100%. **(C)** The levels of glutathione (GSH) and glutathione disulfide (GSSG) were determined in untreated Calu-3 cells using the GSH/GSSG-Glo Assay (Promega) according to the manufacturer’s instructions. The results are from 4 biological replicates with 3 technical replicates for **C**. Error bars represent the standard deviation (SD). *p*-values were calculated using an unpaired two-tailed t-Test.\**p*□<□0.05; \*\**p*□<□0.01.

Previous studies already revealed strong variations in the susceptibilities of different cell lines towards allicin, which were caused by different intracellular GSH contents (Gruhlke et al 2016, Gruhlke et al 2019). Thus, we measured the intracellular GSH and GSSG levels in Vero E6 and Calu-3 cells **(Fig. 1C)**. The GSH content of the more tolerant Calu-3 cells was determined as 3.2 μM, which was 4.2-fold higher compared to only 0.77 μM GSH as measured in Vero E6. As expected, the amounts of GSSG were very low with 0.05 μM and 0.009 μM in Calu-3 and Vero E6 cells, respectively. These data support that Calu-3 cells show greater allicin tolerance due to higher GSH levels compared to Vero E6 cells.

### Allicin inhibits SARS-CoV-2 replication in Vero E6 and Calu-3 cells

The antiviral effect of allicin on SARS-CoV-2 replication in Calu-3 and Vero E6 cells was analyzed. Therefore, different treatment options of infected Vero E6 cells were compared: (1) Cells were pre-exposed to 50 μM allicin for 30 min before SARS-CoV-2 infection. (2) The virus was treated with 50 μM allicin for 30 min prior to infection. (3) Infected Vero E6 cells were exposed to 50 μM allicin for 30 min post infection (p.i.) **(Fig. 2A)**. The number of infectious SARS-CoV-2 particles (PFU, plaque forming units) was determined 24 h p.i. by the plaque titration assay. However, only the addition of 50 μM allicin to infected Vero E6 cells led to a significant 70% decrease in the amount of infectious virus particles, whereas exposure of cells or the virus to allicin prior to infection caused only a 16-21 % reduction of viral plaques **(Fig. 2A)**. These results suggest that allicin affects host-pathogen interactions by antiviral and immunomodulatory activities.

**Fig. 2.**
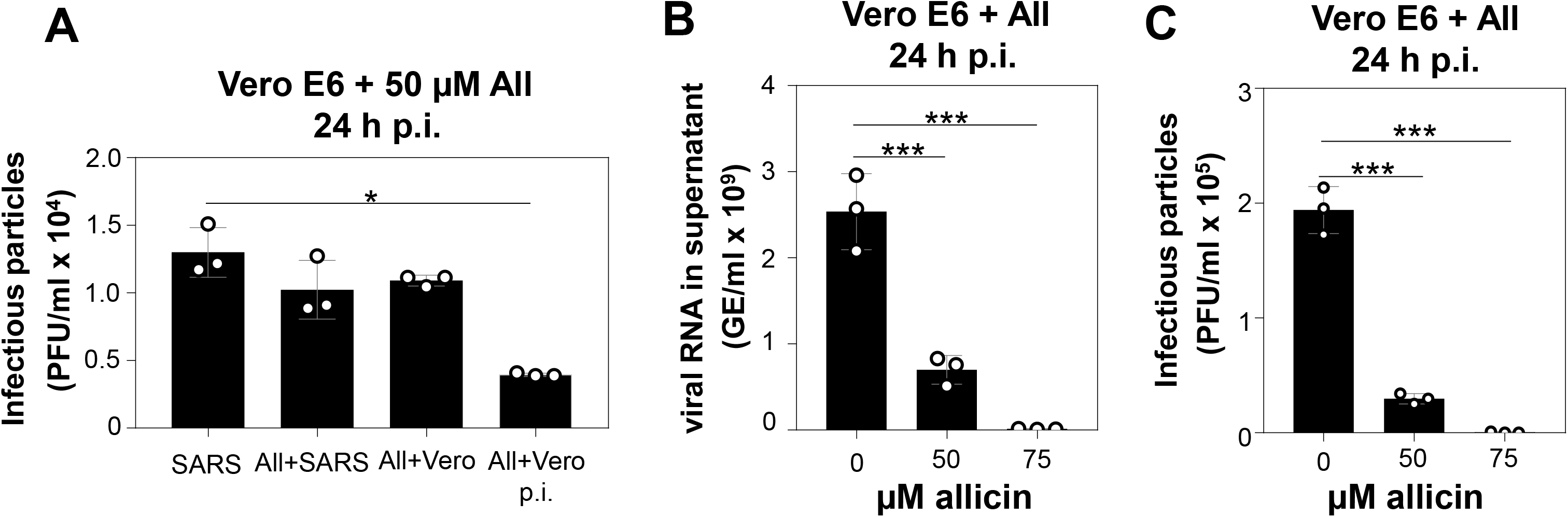
Allicin treatment of SARS-CoV-2 infected Vero E6 cells inhibits viral replication. **(A-C)** Vero E6 cells were infected with SARS-CoV-2 at a MOI of 0.01. After 24 h p.i. viral replication was analyzed by determination of infectious viral particles or viral RNA in the supernatant. **(A)** Comparison of different allicin treatment options: Untreated Vero E6 cells infected with SARS-CoV-2 as control (SARS), SARS-CoV-2 pre-treated with 50 μM allicin for 30 min prior to infection of host cells (All+SARS), host cells pre-treated with 50 μM allicin for 30 min prior to infection with SARS-CoV-2 (All+Vero) and SARS-CoV-2 infected host cells treated with 50 μM allicin p.i. (All+Vero p.i.). Allicin treatment p.i. significantly decreased the number of infectious viral particles. **(B, C)** The amount of viral RNA **(B)** and infectious viral particles **(C)** was determined after treatment of SARS-CoV-2 infected Vero E6 cells with 50 and 75 μM allicin p.i. Increased allicin doses significantly inhibit SARS-CoV-2 replication. The results **A-C** are from 3 biological replicates with 2 technical replicates for **B**. Error bars represent the SD. *p*-values were calculated using an unpaired two-tailed t-Test.\**p*□<□0.05. \*\*\**p*□<□0.001.

We further investigated viral replication after allicin exposure by determination of viral RNA genome equivalents (GE) from the supernatant of infected cells using quantitative RT-PCR. In agreement with the plaque assays, the qRT-PCR results revealed a 72% lower amount of viral RNA after addition of 50 μM allicin to SARS-CoV-2 infected Vero E6 cells **(Fig. 2B, C)**. Moreover, virus plaque assays and qRT-PCR results showed an almost complete >99 % inhibition of SARS-CoV-2 replication after exposure to 75 μM allicin, supporting the strong antiviral activity of allicin in infected Vero E6 cells **(Fig 2B, C)**.

Additionally, the antiviral effects of biocompatible doses of allicin were analyzed in the human lung cell line Calu-3. After infection with SARS-CoV-2 at a multiplicity of infection (MOI) of 0.01 and 0.005 Calu-3 cells were treated with biocompatible doses of 100 and 200 μM allicin and analyzed 16 h p.i. and 24 h p.i., respectively **(Fig. 3)**. Treatment of infected Calu-3 cells with 100 μM allicin did not significantly inhibit viral replication **(Fig. 3A–D)**. However, exposure of infected Calu-3 cells to 200 μM allicin led to a significant >60 % decrease of viral RNA **(Fig. 3A, B)** and a >65% reduction of infectious particles **(Fig. 3C, D)**.

**Fig. 3.**
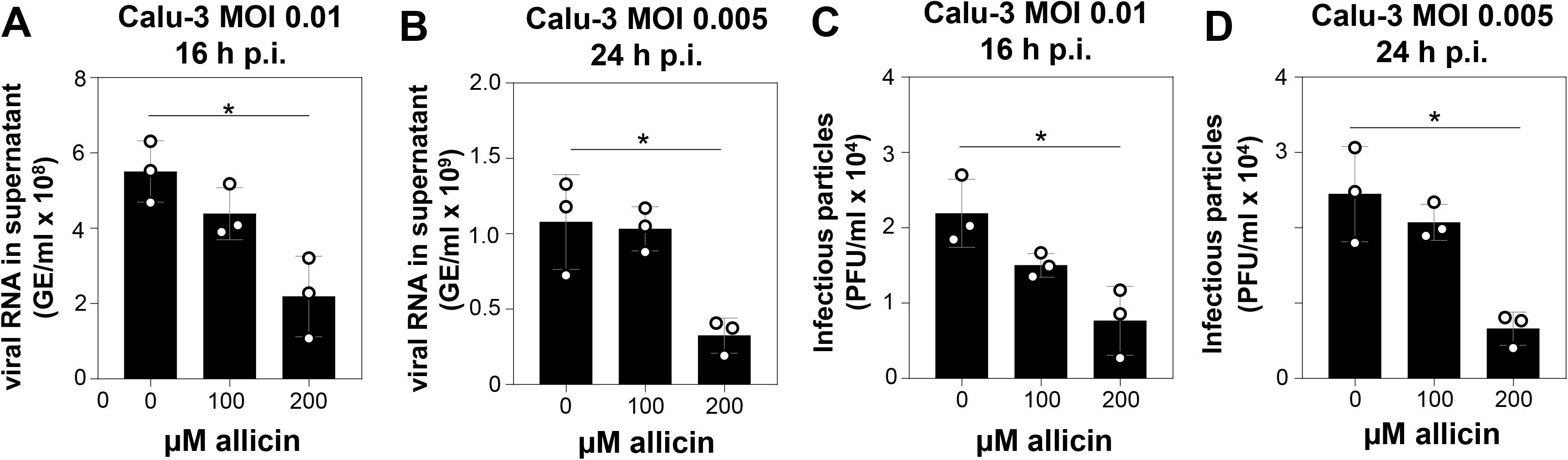
Allicin inhibits SARS-CoV-2 replication in human Calu-3 lung cells as indicated by decreased amounts of viral RNA and infectious particles. **(A-D)** SARS-CoV-2 infected Calu-3 cells were treated with 100 and 200 μM allicin p.i.. The amount of viral RNA **(A, B)** and infectious viral particles **(C, D)** was determined 16 h **(A, C)** and 24 h **(B, D)** p.i. with SARS-CoV-2 at a MOI of 0.01 **(A, C)** and 0.005 **(B, D)**. 200 μM allicin significantly inhibit viral replication after 16 h and 24 h. The results are from 3 biological replicates with 2 technical replicates for **A** and **B**. Error bars represent the SD. *p*-values were calculated using an unpaired two-tailed t-Test. \**p*□<□0.05.

The antiviral effect of allicin on SARS-CoV-2 infected Calu-3 cells was further supported by microscopy imaging **(Fig. 4)**. While SARS-CoV-2 infection at a MOI of 0.01 resulted in cellular damage of Calu-3 cells after 24 h p.i., the addition of allicin partially protected the cells against this damage **(Fig. 4)**. Taken together, our results show that allicin exerts a strong antiviral effect and inhibits SARS-CoV-2 replication in both the primate kidney-derived cell line Vero E6 and the human lung cell line Calu-3.

**Fig. 4.**
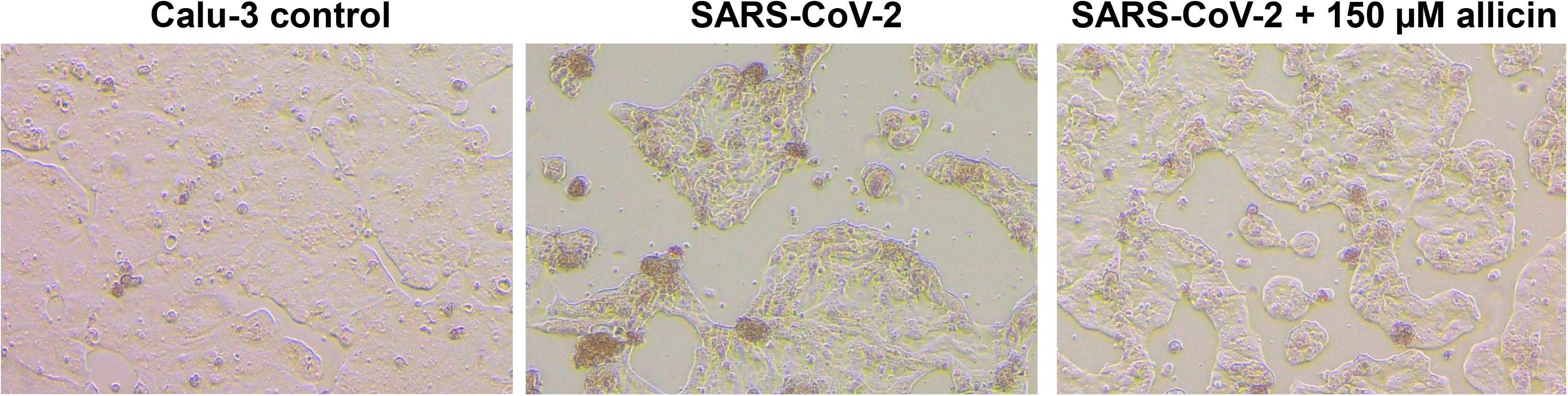
Allicin protects Calu-3 cells against SARS-CoV-2 damage. SARS-CoV-2 induced cellular effects were studied in Calu-3 cells 24 h p.i. Calu-3 cells were infected with SARS-CoV-2 at a MOI of 0.01. At 24 h p.i., Calu-3 cells showed cellular damages, including cell rounding, detachment and cell death. Allicin treatment decreased the observed cellular damage significantly. Cells were imaged with a Nikon Ts2R-FL inverted microscope.

### Changes in the Calu-3 proteome after SARS-CoV-2 infection

Label-free quantitative (LFQ) proteomics by Orbitrap Fusion LC-MS/MS analysis was used to investigate the changes in the proteome of Calu-3 cells after SARS-CoV-2 infection and the effect of allicin. The proteome samples of Calu-3 cells were analyzed before infection (Mock) and 24 h p.i. with SARS-CoV-2 at a MOI of 0.01 in the absence or presence of 150 μM allicin in 3-4 biological and 1-3 technical replicates. The total LFQ intensities of all proteins in each sample were normalized and represent 100% of the total protein abundance. Overall, we quantified 4251 proteins, including 4243 Calu-3 host proteins and 8 SARS-CoV-2 proteins in the total proteome (**Tables S1-S2)**. After 24 h of SARS-CoV-2 infection, about 207 and 329 proteins were ≥1.5-fold induced and <0.66-fold decreased, respectively **(Table S3)**. These 536 differentially expressed proteins contribute only to 2.73 % of the total proteome abundance in SARS-CoV-2 infected Calu-3 cells **(Table S3)**.The proteins were sorted into KEGG Ontology (KO) or Uniprot categories and their fold-changes, *p*-values and averaged abundances were visualized in Voronoi treemaps as color gradients and cell sizes, respectively **(Fig. 5A–D**, **Table S3)**. A subset of the most strongly induced proteins in the Calu-3 proteome after SARS-CoV-2 infection is listed in **Table 1**.

**Fig. 5.**
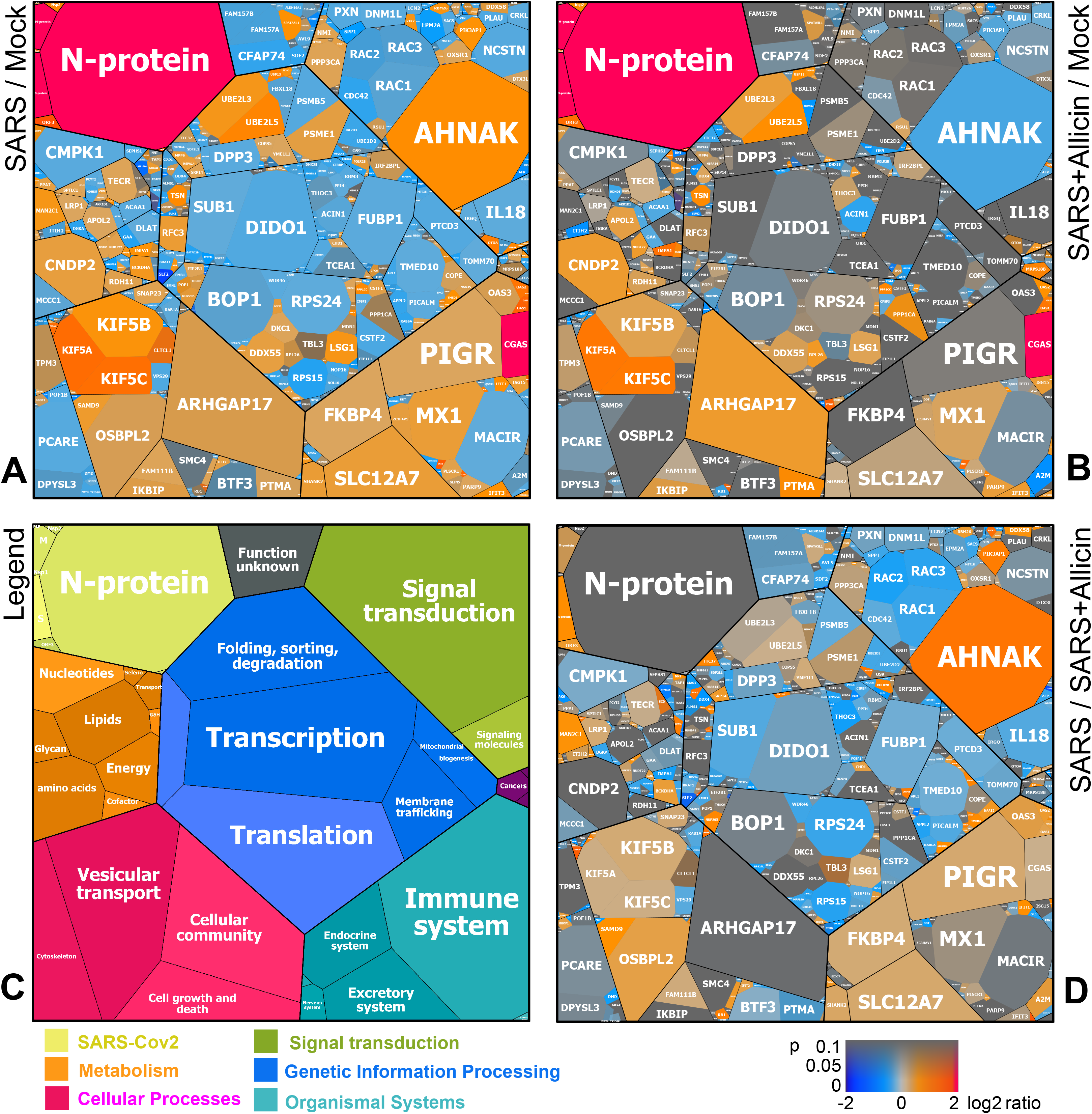
Calu-3 proteome treemaps reveal the protective effect of allicin after SARS-CoV-2 infection. The host-viral proteome treemaps **(A, B, D)** show only the 536 differentially expressed proteins upon SARS-CoV-2 infection and were constructed by the Paver software (Mehlan et al 2013). The treemaps visualize the following proteome changes: **A)** SARS-CoV-2 infection/ Mock, **B)** SARS-CoV-2 infection + Allicin/ Mock and **D)** SARS-CoV-2 infection − / + Allicin. The treemap **(C)** serves as legend for the functional KEGG categories displayed in different colors for level 1 and sublevel 2 as listed in **Table S3**. The cell sizes in **(A, B, D)** denote the average abundances of 207 proteins with ≥1.5-fold inductions and 329 proteins with <0.66-fold decreased expression after SARS-CoV-2 infection. The log2 ratios of the proteins are shown by a red-blue color gradient (red - induction, blue - repression) **(A, B, D)**. *p*-values (*p*□<□0.05; p<0.01) were calculated using an unpaired two-tailed t-Test from 3-4 biological replicates with 1-3 technical replicates **(Table S3)**.

**Table 1.**
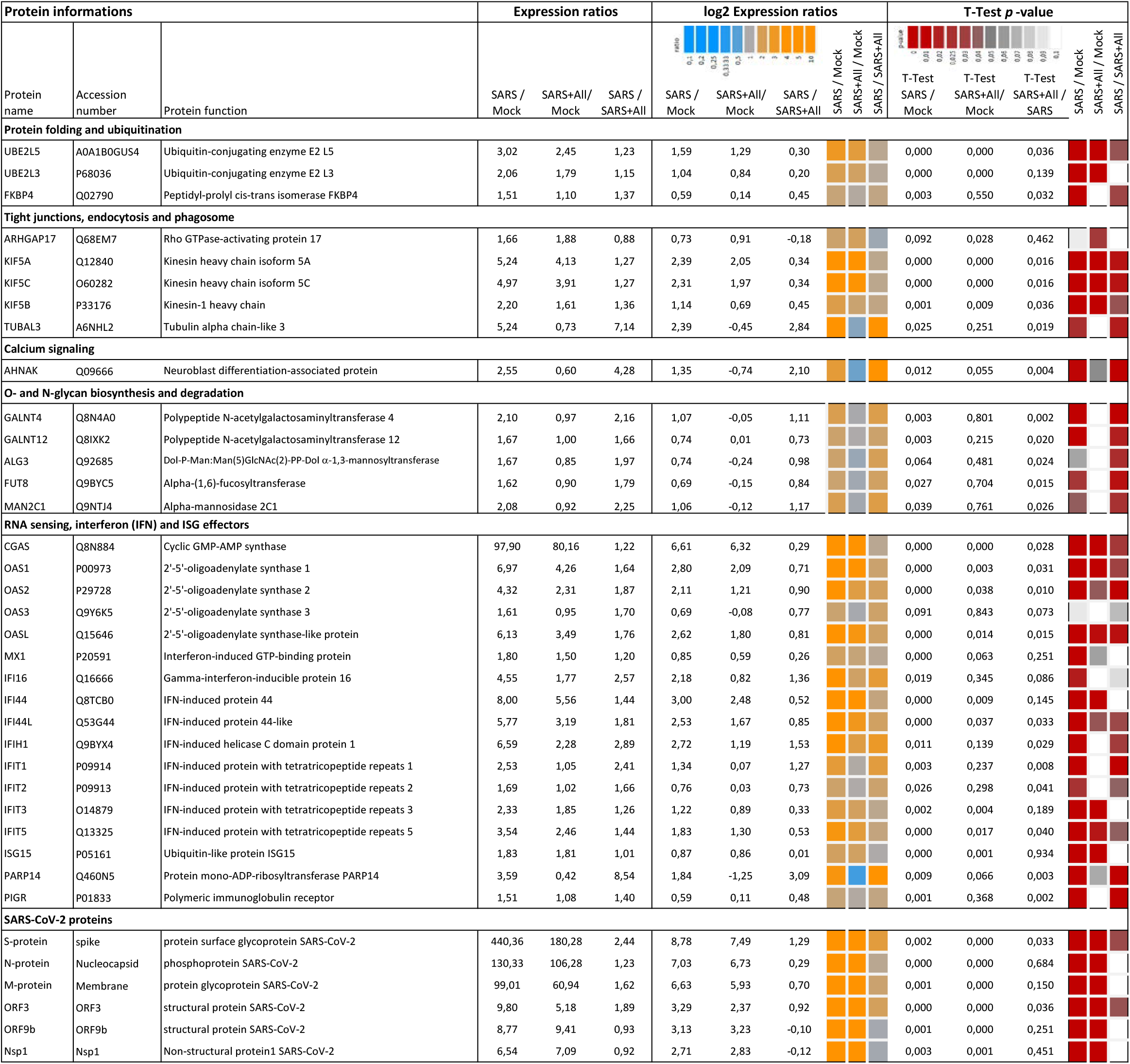
Selected most strongly induced proteins in the Calu-3 proteome after SARS-CoV-2 infection and the effect of allicin treatment p.i. The proteome samples of Calu-3 Mock cells (Mock), SARS-CoV-2 infected Calu-3 cells (SARS) and SARS-CoV-2 infected Calu-3 cells treated with 150 μM allicin p.i. (SARS+All) were harvested after 24 h p.i. and separated by non-reducing SDS PAGE for prefractionation. Protein fractions were tryptic in-gel digested and peptides analyzed by Orbitrap Fusion LC-MS/MS analysis as described in the Methods. The table lists 37 out of 207 identified proteins with >1.5-fold induction upon SARS-CoV-2 infection. These proteins are most strongly induced after SARS-CoV-2 infection, affected by allicin treatment and/or are present at high abundance in the Calu-3 proteome. The proteins were classified according to their KEGG ontologies and UniprotKB annotations. The full set of up- and downregulated proteins after SARS-CoV-2 infection is listed in **Table S3**. The table includes protein names, accession numbers, protein functions, expression ratios, log2 expression ratios and p-values. The log2 ratios and p-values are visualised with a blue-orange and red-grey color code, respectively. p-values were calculated using an unpaired two-tailed t-Test.

| Protein informations |  |  | Expression ratios |  |  | log2 Expression ratios |  |  | T-Test p-value |  |  |  |  |  |  |  |  |
| --- | --- | --- | --- | --- | --- | --- | --- | --- | --- | --- | --- | --- | --- | --- | --- | --- | --- |
| Protein name | Accession number | Protein function | SARS / Mock | SARS+All/ Mock | SARS / SARS+All | SARS / Mock | SARS+All/ Mock | SARS / SARS+All | SARS / Mock | SARS+All / Mock | SARS / SARS+All | T-Test SARS / Mock | T-Test SARS+All/ Mock | T-Test SARS+All / SARS | SARS / Mock | SARS+All / Mock | SARS / SARS+All |
| Protein folding and ubiquitination |  |  |  |  |  |  |  |  |  |  |  |  |  |  |  |  |  |
| UBE2L5 | A0A1B0GUS4 | Ubiquitin-conjugating enzyme E2 L5 | 3,02 | 2,45 | 1,23 | 1,59 | 1,29 | 0,30 |  |  |  | 0,000 | 0,000 | 0,036 |  |  |  |
| UBE2L3 | P68036 | Ubiquitin-conjugating enzyme E2 L3 | 2,06 | 1,79 | 1,15 | 1,04 | 0,84 | 0,20 |  |  |  | 0,000 | 0,000 | 0,139 |  |  |  |
| FKBP4 | Q02790 | Peptidyl-prolyl cis-trans isomerase FKBP4 | 1,51 | 1,10 | 1,37 | 0,59 | 0,14 | 0,45 |  |  |  | 0,003 | 0,550 | 0,032 |  |  |  |
| Tight junctions, endocytosis and phagosome |  |  |  |  |  |  |  |  |  |  |  |  |  |  |  |  |  |
| ARHGAP17 | Q68EM7 | Rho GTPase-activating protein 17 | 1,66 | 1,88 | 0,88 | 0,73 | 0,91 | -0,18 |  |  |  | 0,092 | 0,028 | 0,462 |  |  |  |
| KIF5A | Q12840 | Kinesin heavy chain isoform 5A | 5,24 | 4,13 | 1,27 | 2,39 | 2,05 | 0,34 |  |  |  | 0,000 | 0,000 | 0,016 |  |  |  |
| KIF5C | O60282 | Kinesin heavy chain isoform 5C | 4,97 | 3,91 | 1,27 | 2,31 | 1,97 | 0,34 |  |  |  | 0,000 | 0,000 | 0,016 |  |  |  |
| KIF5B | P33176 | Kinesin-1 heavy chain | 2,20 | 1,61 | 1,36 | 1,14 | 0,69 | 0,45 |  |  |  | 0,001 | 0,009 | 0,036 |  |  |  |
| TUBAL3 | A6NHL2 | Tubulin alpha chain-like 3 | 5,24 | 0,73 | 7,14 | 2,39 | -0,45 | 2,84 |  |  |  | 0,025 | 0,251 | 0,019 |  |  |  |
| Calcium signaling |  |  |  |  |  |  |  |  |  |  |  |  |  |  |  |  |  |
| AHNAK | Q09666 | Neuroblast differentiation-associated protein | 2,55 | 0,60 | 4,28 | 1,35 | -0,74 | 2,10 |  |  |  | 0,012 | 0,055 | 0,004 |  |  |  |
| O- and N-glycan biosynthesis and degradation |  |  |  |  |  |  |  |  |  |  |  |  |  |  |  |  |  |
| GALNT4 | Q8N4A0 | Polypeptide N-acetylgalactosaminyltransferase 4 | 2,10 | 0,97 | 2,16 | 1,07 | -0,05 | 1,11 |  |  |  | 0,003 | 0,801 | 0,002 |  |  |  |
| GALNT12 | Q8IXK2 | Polypeptide N-acetylgalactosaminyltransferase 12 | 1,67 | 1,00 | 1,66 | 0,74 | 0,01 | 0,73 |  |  |  | 0,003 | 0,215 | 0,020 |  |  |  |
| ALG3 | Q92685 | Dol-P-Man:Man(5)GlcNAc(2)-PP-Dol α-1,3-mannosyltransferase | 1,67 | 0,85 | 1,97 | 0,74 | -0,24 | 0,98 |  |  |  | 0,064 | 0,481 | 0,024 |  |  |  |
| FUT8 | Q9BYC5 | Alpha-(1,6)-fucosyltransferase | 1,62 | 0,90 | 1,79 | 0,69 | -0,15 | 0,84 |  |  |  | 0,027 | 0,704 | 0,015 |  |  |  |
| MAN2C1 | Q9NTJ4 | Alpha-mannosidase 2C1 | 2,08 | 0,92 | 2,25 | 1,06 | -0,12 | 1,17 |  |  |  | 0,039 | 0,761 | 0,026 |  |  |  |
| RNA sensing, interferon (IFN) and ISG effectors |  |  |  |  |  |  |  |  |  |  |  |  |  |  |  |  |  |
| CGAS | Q8N884 | Cyclic GMP-AMP synthase | 97,90 | 80,16 | 1,22 | 6,61 | 6,32 | 0,29 |  |  |  | 0,000 | 0,000 | 0,028 |  |  |  |
| OAS1 | P00973 | 2'-5'-oligoadenylate synthase 1 | 6,97 | 4,26 | 1,64 | 2,80 | 2,09 | 0,71 |  |  |  | 0,000 | 0,003 | 0,031 |  |  |  |
| OAS2 | P29728 | 2'-5'-oligoadenylate synthase 2 | 4,32 | 2,31 | 1,87 | 2,11 | 1,21 | 0,90 |  |  |  | 0,000 | 0,038 | 0,010 |  |  |  |
| OAS3 | Q9Y6K5 | 2'-5'-oligoadenylate synthase 3 | 1,61 | 0,95 | 1,70 | 0,69 | -0,08 | 0,77 |  |  |  | 0,091 | 0,843 | 0,073 |  |  |  |
| OASL | Q15646 | 2'-5'-oligoadenylate synthase-like protein | 6,13 | 3,49 | 1,76 | 2,62 | 1,80 | 0,81 |  |  |  | 0,000 | 0,014 | 0,015 |  |  |  |
| MX1 | P20591 | Interferon-induced GTP-binding protein | 1,80 | 1,50 | 1,20 | 0,85 | 0,59 | 0,26 |  |  |  | 0,000 | 0,063 | 0,251 |  |  |  |
| IFI16 | Q16666 | Gamma-interferon-inducible protein 16 | 4,55 | 1,77 | 2,57 | 2,18 | 0,82 | 1,36 |  |  |  | 0,019 | 0,345 | 0,086 |  |  |  |
| IFI44 | Q8TCB0 | IFN-induced protein 44 | 8,00 | 5,56 | 1,44 | 3,00 | 2,48 | 0,52 |  |  |  | 0,000 | 0,009 | 0,145 |  |  |  |
| IFI44L | Q53G44 | IFN-induced protein 44-like | 5,77 | 3,19 | 1,81 | 2,53 | 1,67 | 0,85 |  |  |  | 0,000 | 0,037 | 0,033 |  |  |  |
| IFIH1 | Q9BYX4 | IFN-induced helicase C domain protein 1 | 6,59 | 2,28 | 2,89 | 2,72 | 1,19 | 1,53 |  |  |  | 0,011 | 0,139 | 0,029 |  |  |  |
| IFIT1 | P09914 | IFN-induced protein with tetratricopeptide repeats 1 | 2,53 | 1,05 | 2,41 | 1,34 | 0,07 | 1,27 |  |  |  | 0,003 | 0,237 | 0,008 |  |  |  |
| IFIT2 | P09913 | IFN-induced protein with tetratricopeptide repeats 2 | 1,69 | 1,02 | 1,66 | 0,76 | 0,03 | 0,73 |  |  |  | 0,026 | 0,298 | 0,041 |  |  |  |
| IFIT3 | O14879 | IFN-induced protein with tetratricopeptide repeats 3 | 2,33 | 1,85 | 1,26 | 1,22 | 0,89 | 0,33 |  |  |  | 0,002 | 0,004 | 0,189 |  |  |  |
| IFIT5 | Q13325 | IFN-induced protein with tetratricopeptide repeats 5 | 3,54 | 2,46 | 1,44 | 1,83 | 1,30 | 0,53 |  |  |  | 0,000 | 0,017 | 0,040 |  |  |  |
| ISG15 | P05161 | Ubiquitin-like protein ISG15 | 1,83 | 1,81 | 1,01 | 0,87 | 0,86 | 0,01 |  |  |  | 0,000 | 0,001 | 0,934 |  |  |  |
| PARP14 | Q460N5 | Protein mono-ADP-ribosyltransferase PARP14 | 3,59 | 0,42 | 8,54 | 1,84 | -1,25 | 3,09 |  |  |  | 0,009 | 0,066 | 0,003 |  |  |  |
| PIGR | P01833 | Polymeric immunoglobulin receptor | 1,51 | 1,08 | 1,40 | 0,59 | 0,11 | 0,48 |  |  |  | 0,001 | 0,368 | 0,002 |  |  |  |
| SARS-CoV-2 proteins |  |  |  |  |  |  |  |  |  |  |  |  |  |  |  |  |  |
| S-protein | spike | protein surface glycoprotein SARS-CoV-2 | 440,36 | 180,28 | 2,44 | 8,78 | 7,49 | 1,29 |  |  |  | 0,002 | 0,000 | 0,033 |  |  |  |
| N-protein | Nucleocapsid | phosphoprotein SARS-CoV-2 | 130,33 | 106,28 | 1,23 | 7,03 | 6,73 | 0,29 |  |  |  | 0,000 | 0,000 | 0,684 |  |  |  |
| M-protein | Membrane | protein glycoprotein SARS-CoV-2 | 99,01 | 60,94 | 1,62 | 6,63 | 5,93 | 0,70 |  |  |  | 0,001 | 0,000 | 0,150 |  |  |  |
| ORF3 | ORF3 | structural protein SARS-CoV-2 | 9,80 | 5,18 | 1,89 | 3,29 | 2,37 | 0,92 |  |  |  | 0,000 | 0,000 | 0,036 |  |  |  |
| ORF9b | ORF9b | structural protein SARS-CoV-2 | 8,77 | 9,41 | 0,93 | 3,13 | 3,23 | -0,10 |  |  |  | 0,001 | 0,000 | 0,251 |  |  |  |
| Nsp1 | Nsp1 | Non-structural protein1 SARS-CoV-2 | 6,54 | 7,09 | 0,92 | 2,71 | 2,83 | -0,12 |  |  |  | 0,003 | 0,001 | 0,451 |  |  |  |

The proteome after SARS-CoV-2 infection revealed altered expression of various cellular pathways, including the interferon-stimulated gene (ISG) signature, transcription, translation and protein degradation, the cytoskeleton, vesicular trafficking and tight junctions, apoptosis, signal transduction pathways as well as carbon, lipid and nucleotide metabolism **(Fig. 5A**, **Table 1**, **Table S3)**. In addition, the 8 detected SARS-CoV-2 proteins were induced after 24 h p.i. of Calu-3 cells, with the ribonucleocapsid protein (N-protein) representing with 0.35% of the total proteome one of the most abundant proteins in infected Calu-3 cells. The N-protein was 29- and 21-fold higher expressed compared to the membrane protein (M-protein) (0.012 %) and spike protein (S-protein) (0.016%), respectively **(Fig. 6**, **Table S3)**, confirming previous data with infected Vero E6 cells (Zecha et al 2020). The viral proteins Nsp1, Nsp2, ORF3, ORF9b and the papain-like protease PLP were low abundant, contributing from 0.00022% (PLP) to 0.0043% (ORF3) to the total proteome, while other viral proteins were not detected.

**Fig. 6.**
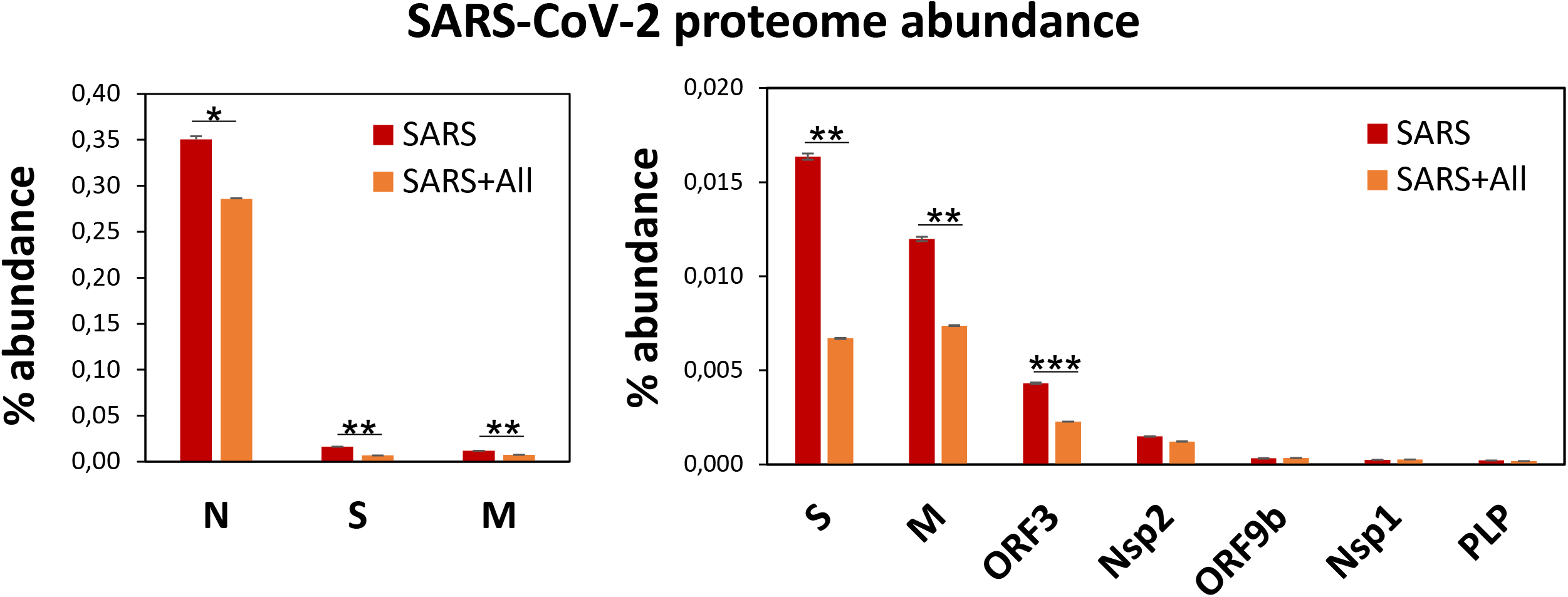
Allicin treatment leads to a decreased abundance of SARS-CoV-2 proteins in the proteome of infected Calu-3 cells. The abundance of the SARS-CoV-2 proteins 24 h p.i. relative to the total proteome abundance of infected Calu-3 cells was calculated in the absence (SARS) or presence of 150 μM allicin exposure p.i. (SARS+All). The N-protein showed a 21-29-fold higher abundance compared to the structural proteins S and M and was omitted in the right panel. The relative abundance of the structural proteins N, S, M and ORF3 was significantly decreased in the proteome of infected Calu-3 cells after allicin treatment. The proteome results were obtained from 3-4 biological replicates with 1-3 technical replicates **(Table S3)**. Error bars represent the SD*. p*-values were calculated using an unpaired two-tailed t-Test. \**p*□<□0.05; **p<0.01; ***p<0.001.

SARS coronaviruses were shown to enter the cell via endocytosis and direct fusion with the cell membrane (Ou et al 2020, Wang et al 2008). In agreement with these reports, 18 proteins involved in vesicular transport and cytoskeleton regulation, such as formation of lysosomes, phagosomes and exosomes were 1.5-5.2-fold higher expressed after infection in the Calu-3 cell proteome **(Fig. 5A**, **Table 1**, **Table S3)**. Among these proteins are the abundant and highly induced kinesins (KIF5A/B/C), clathrin (CLTCL1) and tubulin (TUBAL3), which are microtubule-associated proteins and participate in endocytosis and traffic of viral RNA and vesicles. The 1.7-fold induced highly abundant Rho GTPase-activating protein 17 (ARHGAP17) could be involved in the repair of tight junctions, which are often damaged in COVID-19 patients (De Maio et al 2020, Tian et al 2020).

About 21 proteins of the interferon (IFN) and ISG response were strongly induced, including sensors of viral RNA, the JAK-STAT signal transduction pathway and antiviral effectors that interfere with the viral life cycle **(Fig. 5A**, **Table 1**, **Table S3)** (Schneider et al 2014). For a better understanding, the RNA sensing receptors, IFN and ISG signaling cascades and the previously described antiviral functions of the ISG effectors are displayed in a schematic **(Fig. 7AB)**. The cyclic GMP-AMP (cGAMP) synthase (cGAS) was most strongly 98-fold upregulated upon infection, acting as sensor of viral RNA **(Table 1**, **Table S3**, **Fig. 5A)** (Schneider et al 2014). cGAMP activates the stimulator of interferon genes (STING) **(Fig. 7A)**. The 2’-5’-oligoadenylate synthases (OAS1-3, OASL) were 1.6-7-fold induced upon infection to produce 2’-5’-adenylic acid as second messenger and activator of RNaseL for viral RNA degradation. The IFN-induced helicase C-domain containing protein (IFIH) was 6.5-fold upregulated, which activates the mitochondrial antiviral signaling protein (MAVS) to induce the IFN response. Other IFN-induced effector proteins with tetratricopeptide repeats (IFIT1-3, IFIT5) were 1.6-3.5-fold induced after infection and function in RNA degradation and inhibition of translation. Further effectors are the Interferon-induced myxoma resistance protein 1 (MX1) and the Polymeric immunoglobulin receptor (PIGR), which represented 0.05 and 0.1% of the total proteome abundance and were 1.8 and 1.5-fold induced, respectively. MX1 is a dynamin-like GTPase, which forms ring-like structures and traps incoming ribonucleocapsids, thereby blocking uncoating and vesicular trafficking to direct them for degradation **(Fig. 7B)** (Schneider et al 2014). MX1 was also reported to be up-regulated in COVID-19 patients (Bizzotto et al 2020).

**Fig. 7.**
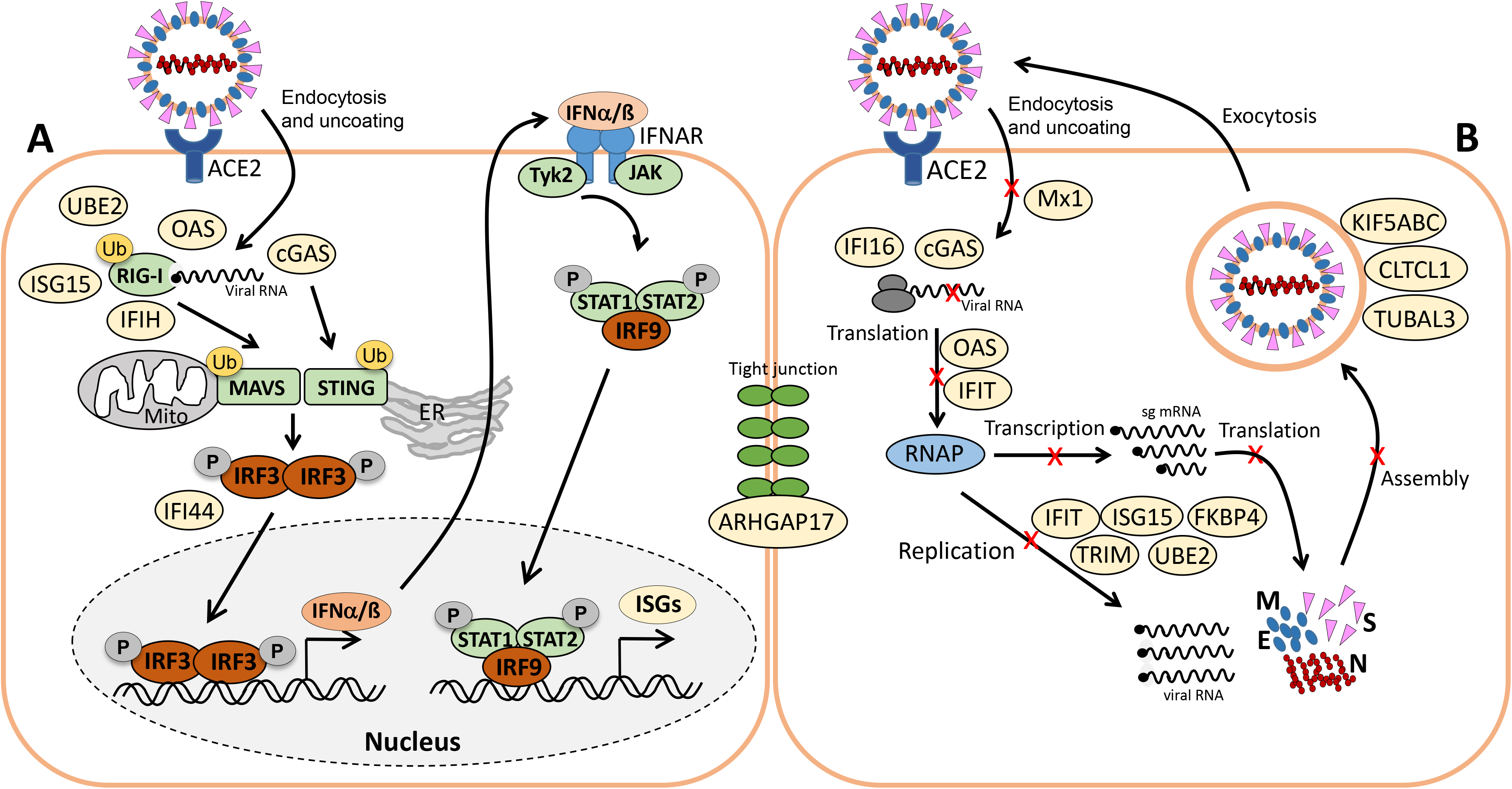
Schematic of viral RNA recognition, activation of the IFN and ISG signaling pathways (A) and antiviral functions of the identified ISG effectors (B). **A)** SARS-CoV-2 enters host cells via endocytosis. RIG-I is a cytosolic receptor to recognize viral RNA. cGAS and OAS are ISG effectors that function as RNA sensors. RIG-I, IFIH and cGAS activate the mitochondrial antiviral-signaling protein (MAVS) and stimulator of IFN genes (STING), leading to phosphorylation of IFN responsive factors (e.g. IRF3), followed by IRF3 dimerization, translocation into the nucleus and transcriptional activation of IFN expression. IRF3 is negatively regulated by IFI44. RIG-I, MAVS and STING can be regulated by ubiquitination (UBE2) and ISGylation (ISG15). Type-I IFN a/ß bind to the IFNAR receptor, resulting in phosphorylation of signal transducers and activators of transcription (STAT1/2) by the JAK and TYK kinases. Phosphorylated STAT1/2 form dimers and bind to IRF9, which triggers transcription of IFN-stimulated genes (ISGs) in the nucleus. **B)** Antiviral ISG effectors affect different stages of the viral life cycle. Mx1 inhibits virus endocytosis and uncoating of the ribonucleocapsid. IFI16, OAS and IFIT function in viral RNA degradation and block translation. IFIT, ISG15, TRIM and UBEL inhibit transcription, replication, translation or virus assembly. FKBP4 promotes protein folding. Kinesins (KIFA/B/C), Clathrin (CLTCL1) and TUBAL3 are involved in transport of virus vesicles. ARHGAP17 facilitates the formation or repair of tight junctions. The figure is adapted from reference (Schneider et al 2014).

Furthermore, the abundant cytokine IL18, the IL-1 receptor antagonist protein (IL1RN), the macrophage immunometabolism regulator MACIR and the Alpha-2-macroglobulin (A2M) were ~0.6-fold lower expressed in infected cells. MACIR is implicated in the regulation of macrophages and autoimmune diseases (McGauran et al 2020). Alpha-2-macroglobulin accounts for approximately 10% of the serum antiprotease capacity and was shown to inhibit SARS-CoV-2 entry (Oguntuyo et al 2020).

An important role in signal transduction and regulation of the antiviral response plays the abundant ISG15 effector, which was 1.8-fold induced after SARS-CoV-2 infection in the proteome. ISG15 functions amongst others as Ubiquitin-like modifier in ISGylation of RIG-I and IRF-3, which are targeted for degradation or activated to regulate IFN and ISG production **(Fig. 7A)** (Masucci 2020). Widespread ISGylation of newly synthesized viral proteins is proposed to inhibit viral replication and translation (Durfee et al 2010).

Additionally, post-translational modification by polyubiquitination of host signaling factors, such as RIG-I, STING and MAVS is important for regulation of the IFN response upon SARS-CoV-2 infection **(Fig. 7A)**. Thus, several ubiquitin-conjugating E2 enzymes (UBE2L3, UBE2L5), the E3 ubiquitin ligases (TRIM21, TRIM38, ARIH2) and the ubiquitin specific protease or deconjugases (USP13) are 1.5-3.2-fold induced in the infected cells, while other E2, E3 enzymes and deconjugases (e.g. UBE2D2/3, RNF214, USP4, USP47, USP48) are 0.2-0.62-fold lower expressed **(Table 1**, **Table S3**, **Fig. 5A)**. Since host and viral targets of ubiquitination and ISGylation are often directed to degradation, components of the proteasome, proteases, protein folding factors, and chaperones are 1.5-1.8-fold upregulated. The folding factors include the highly abundant peptidyl-prolyl cis-trans isomerase FKBP4, which functions as immunophilin and co-chaperone to interact with HSP90.

Apart from protein modification, the virus relies on protein synthesis and translation by the host machinery for its successful replication and infectivity. Accordingly, 24 proteins involved in translation were 1.5-2.8-fold upregulated under SARS-CoV-2 infection, including translation factors EIF2B1, ribosomal proteins (RPL26, MRPS30, RRP8, PDCD11, MRPL4), RNA helicases (DDX55, DDX56), RNAses (POP1, XRN1) and other regulatory factors, such as phosphatases (PPP1CC, PPP1CA, PPP2R5A) **(Table S3**, **Fig. 5A)**.

In addition, 16 proteins involved in transcription and the spliceosome were upregulated in infected Calu-3 cells, including the pre-mRNA splicing factors Slu7, PRPF40B, SCAF11 and the U1 small nuclear ribonucleoprotein C (SNRPC), which were 1.6-1.8-fold higher expressed. The transcription factors GABPA, ZNF579, SP110 and TSC22D2 were also induced after infection. However, the majority of differentially expressed proteins involved in transcription (48) and translation (30) were repressed after SARS-CoV-2 infection, including the highly abundant proteins DIDO1, SUB1, FUBP1, TCEA1, BOP1, RPS24 and RPS15 **(Table S3**, **Fig. 5A)**.

Moreover, virus replication and proliferation inside host cells requires reprogramming of the host metabolism, which was evident by the upregulation of 34 proteins and downregulation of 43 proteins involved mainly in lipid, energy, glycan and nucleotide metabolism **(Table 1**, **Table S3**, **Fig. 5A)**. The induced proteins might function in the biosynthesis of the building blocks for viral phospholipid membranes, glycosylation of surface proteins and viral RNA genomes. Since the nucleotide pool is essential for coronavirus replication (Bojkova et al 2020), some purine and pyrimidine biosynthesis proteins were 1.7-2.3-fold induced (NT5C2, UPP1, PPAT), while others were 0.5-0.65-fold repressed (CMPK1, AK6, ENPP4) **(Table S3**, **Fig. 5A)**.

Furthermore, expression of several signaling pathways, including JAK-STAT, MAPK, Wnt, Ras and Rap1 signaling were affected by SARS-CoV-2 infection. The JAK-STAT pathways senses and transduces IFN-signals via a phosphorylation cascade to activate ISG expression **(Fig. 7A)**. Thus, STAT2, N-myc interactor NMI and the RIG-I receptor were 1.6-1.8-fold induced upon infection **(Table S3**, **Fig. 5A)**. Proteins of the MAPK signaling pathways were activated in response to infections with SARS-CoV (Bouhaddou et al 2020) and 1.6-1.8-fold induced in the proteome of SARS-CoV-2 infected cells. Proteins of the PI3K/Akt signaling pathway were 2.3-2.6-fold upregulated in infected cells, controlling apoptosis of host cells for successful viral replication. The highly abundant neuroblast differentiation-associated protein AHNAK was 2.6-fold induced after SARS-CoV-2 infection. AHNAK is required for calcium signaling and might regulate the immune response (Matza et al 2009). Proteins of the Ras-signaling pathway were 0.5-0.64-fold downregulated upon virus infection, including three Rac GTPases Rac1-3 that are implicated in the regulation of cell morphology, migration and invasion, by transducing signals from cell surface receptors to the actin and microtubule cytoskeletons (Wheeler et al 2006). Similarly, other proteins involved in the cytoskeleton organisation were 0.3-0.66-fold lower expressed, indicating re-organization of the cytoskeleton for transport of virus particles.

### Allicin leads to a decreased antiviral IFN response in the proteome of infected Calu-3 cells

Next, we investigated the effect of allicin on the proteome changes upon SARS-CoV-2 infection. Quantification of the 8 viral proteins in infected Calu-3 cells after allicin treatment revealed a significantly 18-59% decreased abundance of the structural proteins N, M and S and ORF3, supporting the antiviral effect of allicin in the proteome **(Fig. 6)**.

Allicin treatment resulted in a diminished IFN-response in SARS-CoV-2 infected cells, since expression of innate immune receptors and ISG effectors of the JAK-STAT signaling pathways were strongly decreased, including FKBP4, PIGR, MX1, cGAS, OAS1-3 and IFIT1-3 **(Table 1**, **Table S3**, **Fig. 5B,D)**. In addition, proteins involved in ubiquitination (UBE2L3/5) and the JAK/STAT, MAPK, PI3K/Akt and Ras signaling pathways showed lower expression changes after allicin treatment. The abundant calcium-signaling protein AHNAK was repressed after allicin exposure, while it was induced in infected cells. Allicin resulted in decreased expression of kinesins KIFA/B/C, clathrin CLTCL1 and tubulin TUBAL3, indicating reduced endocytosis and traffic of vesicles. Moreover, prothymosin alpha (PTMA), which confers resistance to infections, was 2.6-fold upregulated after allicin exposure.

Similarly, the expression of proteins involved in transcription, spliceosome and translation was restored to the levels of uninfected (Mock) cells after allicin exposure of SARS-CoV-2 infected cells, including the abundant proteins DIDO1, SUB1, FUBP1, TCEA1, BOP1, RPS24 and RPS15. Finally, expression of metabolic enzymes involved in glycan, nucleotide and lipid metabolism was normalized to Mock levels by allicin, including GALNT4/12, ALG3, FUT8, MAN2C1, CMPK1 and TECR **(Table 1**, **Table S3**, **Fig. 5B,D)**. Overall, allicin showed antiviral effects in the host proteome as revealed by the diminished IFN-dependent antiviral response, the effect on signal transduction, transcription, translation and metabolism, which was reversed to Mock levels.

Finally, we monitored *S*-thioallylations in the proteome of SARS-CoV-2 infected Calu-3 cells after 24 h of exposure to 150 μM allicin to identify possible allicin targets. In total, 100 proteins with *S*-thioallylated peptides were identified based on the mass shift of 72 Da at Cys residues **(Table S4)**. These *S*-thioallylated proteins are involved in transcription, translation, ubiquitination and DNA maintenance (31), signal transduction pathways (21), the immune defense (7), lipid, sugar and nucleotide metabolism (8), endocytosis and apoptosis (7), human diseases (4) and unknown functions (10) **(Table S4)**. Interestingly, among the transcription factors are several C2H2 zinc finger proteins (ZNF541, ZNF518A, ZNF33A, ZNF443), containing multiple Cys2-His2 motifs in their DNA binding domains. In addition, some ubiquitin E3 ligases were found *S*-thioallylated (RNF8, HERC4, MEX3C, G2E3), which harbor active site Cys residues (HERC4) or zinc finger domains (RNF8, MEX3C, G2E3). Furthermore, peroxiredoxin-6 (PRDX6), 3-hydroxyacyl-CoA dehydrogenase type-2 (HSD17B10) and cytoplasmic dynein 1 heavy chain 1 (DYNC1H1) proteins were previously found *S*-thioallylated in human Jurkat cells.

## Discussion

Garlic organosulfur compounds, such as allicin were previously shown to act as antivirals against several enveloped viruses, including herpes simplex, parainfluenza, vaccinia and rhinovirus(Rouf et al 2020, Weber et al 1992). These virucidal effects of garlic compounds were proposed to depend on the disruption of the viral envelope and inhibition of viral replication (Rouf et al 2020, Weber et al 1992). Additionally, experiments in pigs showed an immune-enhancing effect of garlic to prevent viral infections by increasing CD8^+^ T- and B-cells, anti-inflammatory cytokines and antibody titres (Guo et al 1993). The various anti-inflammatory and immunomodulatory effects of garlic compounds to improve innate and adaptive immunity are summarized in a recent review (Donma & Donma 2020). In this work, we explored the antiviral effect of allicin on SARS-CoV-2 infected Vero E6 and Calu-3 cells. By determining decreased levels of viral RNA and infectious viral particles, the antiviral effect of allicin against SARS-CoV-2 was demonstrated in both cell lines. However, Calu-3 cells showed a greater allicin tolerance compared to the more sensitive Vero E6 cells. Different sensitivities towards allicin were previously found in various cell lines, such as Jurkat, murine EL-4 T-cells, human lung epithelium carcinoma A549 and human umbilical vein endothelial cells (HUVEC) (Gruhlke et al 2016, Gruhlke et al 2019). The different allicin susceptibilities in these cell lines were caused by different intracellular GSH levels (Gruhlke et al 2016, Gruhlke et al 2019). Allicin leads to *S*-thioallylation of GSH and the formation of S-allylmercaptoglutathione (GSSA), which is accompanied by GSH depletion and an oxidative shift in the GSH redox potential (Gruhlke et al 2010, Gruhlke et al 2019, Müller et al 2016). The measurement of GSH levels confirmed that Calu-3 cells have 4.2-fold higher GSH levels compared to the allicin-sensitive Vero E6 cells.

The significant reduction in viral plaques and RNA copies was evident in both SARS-CoV-2 infected Vero E6 and Calu-3 cells after exposure to sub-lethal doses allicin p.i.. Interestingly, allicin was most effective during host-pathogen interactions to reduce virus replication. Thus, our *in vitro* results support previous studies on the antiviral effect of allicin against other viruses (Rouf et al 2020, Weber et al 1992).

To better understand the antiviral effect of allicin, we investigated the proteome changes of Calu-3 cells upon SARS-CoV-2 infection and the impact of allicin on the host-virus proteome. In agreement with previous proteome studies, SARS-CoV-2 reprograms major host pathways, including signaling pathways, transcription, splicing, translation, protein modification and folding, lipid, glycan and nucleotide metabolism **(Table 1**, **Table S3**, **Fig. 5A**, **Fig. 7AB)** (Bojkova et al 2020, Bouhaddou et al 2020, Zecha et al 2020). The ribonucleocapsid protein was the most abundant viral protein in the infected cell, indicating that a large portion of the translation capacity goes to the N-protein for package of the viral RNA genome.

In addition, our proteome data highlight the importance of the IFN pathway and ISG effectors to prevent virus replication by interacting with various stages of the viral life cycle. Antiviral ISG effectors were among the most highly induced and abundant proteins in the infected host cells, such as MX1, cGAS, OAS1-3, IFIT1-3, ISG15, FKBP4, PIGR and UBE2L3/5, which function in sensing and degradation of viral RNA, inhibition of ribonucleocapsid uncoating, translation and promote the innate immune response **(Table 1**, **Table S3**, **Fig. 5A**, **Fig. 7AB)**. Apart from IFN signaling, proteins involved in motility, tight junction and membrane trafficking are highly induced host proteins, supporting the importance of vesicular transport for virus endocytosis and exocytosis. Thus, our proteomics studies reflect all described host pathways known to be altered after viral infections, suggesting new host targets for SARS-CoV-2 interventions.

At the same time, the proteomic profiling gave the opportunity to monitor the molecular responses of infected Calu-3 cells after allicin treatment. The proteome results of allicin-treated SARS-CoV-2 infected host cells further support the antiviral activity of allicin as revealed by a 18-59% reduced abundance of the structural proteins N, M, S and ORF3. Most expression changes dedicated to virus proliferation are reversed to Mock levels in allicin-treated cells. Allicin strongly affected virus-responsive expression of JAK/STAT, MAPK, PI3K/Akt and Ras signaling pathways, IFN and ISG effectors, transcription, splicing, translation, ubiquitination, vesicular transport, tight junctions as well as glycan, lipid and nucleotide metabolism. This diminished host response is visualized in the treemaps **(Fig. 5A, D)** since the expression profile of SARS versus Mock resembles that of SARS versus SARS+allicin. Thus, our results confirm the antiviral and protective effect of allicin in host cells, supported by a decreased cellular damage of allicin-treated infected Calu-3 cells.

The mode of action of allicin involves *S*-thioallylation of proteins and low molecular weight thiols in bacterial and human Jurkat cells (Gruhlke et al 2019, Loi et al 2019). The majority of *S*-thioallylated Jurkat proteins were abundant cellular proteins, involved in the cytoskeleton, translation and protein folding, although also low abundant redox-sensitive transcription factors, such as MgrA, SarZ, OhrR, HypR, YodB were targets for allicin modification in *S. aureus* and *B. subtilis* cells (Chi et al 2019, Gruhlke et al 2019, Loi et al 2019). In this study, 100 *S*-thioallylated Cys proteins could be verified by LC-MS/MS analysis that are modified by allicin even 24 h post allicin treatment. While the overlap of *S*-thioallylated Cys peptides with previous results in Jurkat cells was low, interesting allicin targets could be identified as C2H2 zinc finger transcription factors (ZNF541, ZNF 518A, ZNF33A, ZNF443) and zinc finger ubiquitin E3 ligases (RNF8, HERC4, MEX3C, G2E3), that impact gene expression. *S*-thioallylation of the Cys residues in the active site and zinc finger domains might lead to Zn^2+^ release and inactivation of these transcription factors and E3 ligases. Allicin treatment further enhanced the IL-1ß-induced production of IL-2 in murine EL-4 T-cells possibly via Zn^2+^ levels (Gruhlke et al 2019). However, it remains to be investigated in future studies whether the immunomodulatory effect of allicin on cytokine secretion in cell cultures is mediated by elevated Zn^2+^ levels or regulation of specific allicin targets by *S*-thioallylation.

Interestingly, no viral *S*-thioallylated Cys peptides were identified, although the spike protein is a Cys-rich glycoprotein exposed on the surface of the virus envelope (Zhang et al 2020). Most likely, human cells have reduced allicin and the majority of *S*-thioallylations within 24 h. Efficient allicin detoxification and removal of *S*-thioallylations was confirmed in yeast and bacterial cells, as shown by fast recovery of growth after a short allicin-induced lag phase (Gruhlke et al 2010, Gruhlke et al 2019, Loi et al 2019, Müller et al 2016).

Taken together, our results support that allicin functions as an antiviral compound that inhibits SARS-CoV-2 replication in two infected cell lines, Vero E6 and Calu-3. However, over-dosage of allicin can be also toxic to human cells, as confirmed in this work. Consequently, the question arises about the biocompatible doses of allicin in humans to prevent bacterial and viral infections (Borlinghaus et al 2021). Allicin is very instable and quickly decomposes to polysulfanes, ajoene and other sulfur compounds during cooking (Block 2010, Borlinghaus et al 2021). In the acidic stomach, the majority of allicin is degraded to 2-propenethiol and allyl methyl sulfide, which are excreted (Block 2010, Borlinghaus et al 2021). In the blood, the effective doses of allicin is further reduced by its reaction with GSH (Block 2010, Borlinghaus et al 2021). Thus, biocompatible and therapeutically relevant concentrations of allicin for the treatment of respiratory tract infections might not be reached with the consumption of garlic-containing food (Borlinghaus et al 2021). Therefore, future investigations should be directed to exploit the volatile nature of biocompatible doses of allicin for the treatment of COVID-19 patients via the pulmonary route to efficiently reach the virus, without damaging host cells.

## Materials and Methods

### Cultivation of cell lines and infection experiments with SARS-CoV-2

Vero E6 (ATCC CRL-1586) and Calu-3 (ATCC HTB-55) cell lines were cultivated in Dulbecco’s Modified Eaglés Medium (DMEM), supplemented with 10% fetal bovine serum (FBS), 1% non-essential amino acids and 1% sodium pyruvate (Gibco), and grown at 37°C and 5% CO_2_ as described previously (Hoffmann et al 2020). Cell lines were free of mycoplasma, authenticated based on morphology and growth properties and confirmed by PCR.

The SARS-CoV-2 Munich isolate (CSpecVir985) was used for infection assays of Vero E6 and Calu-3 cells, under biosafety level 3 with appropriate personal protection. Vero E6 and Calu-3 cells were seeded at densities of 3.5 × 10^5^ cells/ml or 6 × 10^5^ cells/ml, respectively. After 24□h, cells were infected at a MOI of 0.01 or 0.005, diluted in serum-free OptiPro medium for 1 h at 37°C. The medium was removed and cells were washed twice with PBS followed by addition of DMEM and supplements. Cells were harvested 16 and 24 hp.i. For allicin treatment prior infection, host cells or SARS-CoV-2 were pre-incubated with 50 μM allicin for 30 min. Allicin was synthesized by oxidation of 3-[(Prop-2-en-1-yl)disulfanyl]prop-1-ene (diallyl disulfide, Sigma-Aldrich, Germany) with peracetic-acid (glacial acetic acid/H_2_O_2_) as described previously (Gruhlke et al 2010).

### Cell viability assay

The cell viability of Vero E6 and Calu-3 cells was analyzed by quantification of ATP levels using the CellTiter-Glo™ Luminescent Cell Viability assay (Promega) according to the instructions of the manufacturer. The cells were cultivated as described above and exposed to different amounts of allicin for 24 h. For normalization, the cell viability of non-treated cells was set to 100% and the relative viability of cells exposed to allicin was calculated.

### Determination of the levels of GSH and glutathione disulfide (GSSG) in Vero E6 and Calu-3 cells

Vero E6 and Calu-3 cells were cultivated as described above and seeded at densities of 1×10^4^ cells/well. After washing with phosphate-buffered saline, the intracellular GSH and GSSG concentrations were determined using the GSH/GSSG-Glo™ assay (Promega) according to the instructions of the manufacturer for adherent cells. Briefly, total GSH levels were measured in one sample by reduction of GSSG to GSH using DTT. Total GSSG amounts were measured in a second sample by blocking reduced GSH with N-ethylmaleimide (NEM), followed by GSSG reduction with DTT. The GSH transferase (GST) uses GSH as cofactor to convert luciferin-NT to GSH-NT resulting in the release luciferin. Luciferin is oxidized to oxyluciferin by the Ultra-Glo™ rLuciferase, leading to emission of chemiluminescence, which was measured using an integration time of 1 sec/well by the CLARIOstar microplate reader (BMG Labtech). GSH levels were calculated based on GSH standard curves. For determination of the cellular GSH levels, the GSSG amounts were subtracted from the total GSH level.

### Plaque titration assay

The number of infectious virus particles was determined by a plaque titration assay as described previously (Hoffmann et al 2020). Briefly, Calu-3 and Vero E6 monolayers were seeded in 24-well plates and infected with 200 μl of serial dilutions of SARS-CoV-2 containing cell culture supernatants, which were diluted in OptiPro serum-free medium. After 1 h adsorption, the supernatant was removed and cells overlaid with 1.2% Avicel (FMC BioPolymers) diluted in DMEM. After 72 h, the overlay was removed, cells were fixed in 6% formaldehyde and plaques were visualized by crystal violet staining.

### Viral RNA extraction and real-time reverse-transcription PCR

Culture supernatants were used for viral RNA extraction as described previously (Hoffmann et al 2020). RNA extraction was performed from 50 μl culture supernatant with the viral RNA kit (Macherey-Nagel) according to the instructions of the manufacturer. SARS-CoV-2 genome equivalents (GE) were detected by quantitative RT–PCR (LightCycler 480 Real-Time PCR System and Software version 1.5 (Roche)), targeting the SARS-CoV-2 *E* gene and absolute quantification was performed using SARS-CoV-2 specific *in vitro*-transcribed RNA standards (Corman et al 2020).

### Proteome analysis of SARS-CoV-2 infected host cells using Orbitrap Fusion mass spectrometry

6×10^5^ Calu-3 cells per sample were infected with SARS-CoV-2 as described above and treated with 150 μM allicin for 24 h. Calu-3 cells were harvested by centrifugation, the pellets were washed with PBS and alkylated under denaturing conditions in 200 μl urea/ iodoacetamide/ EDTA (UCE-IAM) buffer for 15 min at RT as described (Rossius et al 2018). Subsequently, the alkylated protein extracts were precipitated with trizol and 96% ethanol and washed four times with 1 ml 70% ethanol. The protein pellets were separated by a short 15% non-reducing SDS-PAGE, which was running for 15 min and stained with Colloidal Coomassie Blue. The gel fractions were cut and in-gel tryptic digested as described previously (Chi et al 2011). The eluted peptides were desalted using ZipTip-μC18 material (Merck Millipore) and dissolved in 0.1% (v/v) formic acid before LC-MS/MS analysis. The peptide samples of non-infected Calu-3 cells (Mock) and SARS-CoV-2 infected Calu-3 cells with and without allicin treatment were subjected to nLC-MS/MS analysis using an Orbitrap Fusion (Thermo Fisher Scientific) coupled to a TriVersa NanoMate (Advion, Ltd.) as described previously (Kublik et al 2016). Peptide identification of the human and SARS-CoV-2 proteome was performed by Proteome Discoverer (version 2.2, Thermo Fisher Scientific) using the SequestHT search engine as described (Seidel et al 2018). Human and SARS-CoV-2 proteins were identified by searching all tandem MS/MS spectra against the human proteome protein sequence database (20286 entries) extracted from UniprotKB release 12.7 (UniProt Consortium, Nucleic acids research 2007, 35, D193-197) as well as against the European Virus Archive Global # 026V-03883 sequence database. Peptides were considered to be identified with high confidence at a target false discovery rate of ≤0.01 and with a medium confidence at ≤0.05, based on the q-values. Identified proteins were quantified by the ‘Percursor Ions Quantifier’ implemented in Proteome Discoverer 2.2 based on peak intensities to estimate the abundance of the human and SARS-CoV-2 proteins in the peptide samples. Error tolerance for precursor ion and fragment ion *m/z* values was set to 3 ppm and 0.5 Da, respectively. Two missed cleavage sites were allowed. Methionine oxidation (+15.994915 Da), cysteine carbamidomethylation (+57.021464 Da) and cysteine *S*-thioallylation by allicin (+72.00337 Da for C3H5S1) were set as variable modifications. The mass spectrometry data have been deposited to the ProteomeXchange Consortium via the PRIDE partner repository (Deutsch et al 2020, Perez-Riverol et al 2019) with the dataset identifier PXD024375.

## Data availability

The mass spectrometry data have been deposited to the ProteomeXchange Consortium via the PRIDE partner repository (Deutsch et al 2020, Perez-Riverol et al 2019) with the dataset identifier PXD024375.

## Acknowledgements

This work was supported by an European Research Council (ERC) Consolidator grant (GA 615585) MYCOTHIOLOME and grants from the Deutsche Forschungsgemeinschaft, Germany (AN746/4-1 and AN746/4-2) within the SPP1710 on “Thiol-based Redox switches”, by the SFB973 (project C08) and TR84 (project B06) to H.A. Infections experiments were supported by the TR84 (project A07) to D.N. Mass spectrometry was performed by L.A. at the Centre for Chemical Microscopy (ProVIS) at the Helmholtz Centre for Environmental Research, which is supported by European regional development funds (EFRE-Europe Funds Saxony) and the Helmholtz Association. Support for allicin synthesis was provided by internal funding from the RWTH Aachen University to M.C.H.G. and A.J.S.

## Conflict of Interest

The authors declare to have no conflict of interest.

## Author Contributions

K.M. performed the infection experiments and analyzed the data. V.N.F. and L.A. measured and analyzed the proteomic data. J.B. constructed the Voronoi treemaps. M.C.H.G. and A. S. synthesized allicin. D.N. supervised the infection experiments and gave critical advise. V.N.F. and H.A. conducted the study and wrote the initial manuscript. All authors contributed to the final manuscript.

## Notes

### Competing Interest Statement

The authors have declared no competing interest.

## REFERENCES

Arbach M, Santana TM, Moxham H, Tinson R, Anwar A, Groom M, Hamilton CJ. 2019. Antimicrobial garlic-derived diallyl polysulfanes: Interactions with biological thiols in *Bacillus subtilis*. Biochim Biophys Acta Gen Subj. 1863(6):1050–1058. doi:10.1016/j.bbagen.2019.03.012

Bautista DM, Movahed P, Hinman A, Axelsson HE, Sterner O, Hogestatt ED, Julius D, Jordt SE, Zygmunt PM. 2005. Pungent products from garlic activate the sensory ion channel TRPA1. Proc Natl Acad Sci U S A. 102(34):12248–12252. doi:10.1073/pnas.0505356102

Bizzotto J, Sanchis P, Abbate M, Lage-Vickers S, Lavignolle R, Toro A, Olszevicki S, Sabater A, Cascardo F, Vazquez E, et al. 2020. SARS-CoV-2 Infection Boosts MX1 Antiviral Effector in COVID-19 Patients. iScience. 23(10):101585. doi:10.1016/j.isci.2020.101585

Block E. 2010. Garlic and the other Alliums. The Lore and the Science. RSC Publishing Cambridge UK.

Bojkova D, Klann K, Koch B, Widera M, Krause D, Ciesek S, Cinatl J, Munch C. 2020. Proteomics of SARS-CoV-2-infected host cells reveals therapy targets. Nature. 583(7816):469–472. doi:10.1038/s41586-020-2332-7

Borlinghaus J, Albrecht F, Gruhlke MC, Nwachukwu ID, Slusarenko AJ. 2014. Allicin: chemistry and biological properties. Molecules. 19(8):12591–12618. doi:10.3390/molecules190812591

Borlinghaus J, Foerster J, Kappler U, Antelmann H, Noll U, Gruhlke M, Slusarenko AJ. 2021. Allicin, the odor of freshly crushed garlic: A review of recent progress in understanding allicin’s effects on cells. Molecules. 26(6):1505. doi:https://doi.org/10.3390/molecules26061505

Bouhaddou M, Memon D, Meyer B, White KM, Rezelj VV, Correa Marrero M, Polacco BJ, Melnyk JE, Ulferts S, Kaake RM, et al. 2020. The Global Phosphorylation Landscape of SARS-CoV-2 Infection. Cell. 182(3):685–712 e619. doi:10.1016/j.cell.2020.06.034

Chi BK, Gronau K, Mäder U, Hessling B, Becher D, Antelmann H. 2011. *S*-bacillithiolation protects against hypochlorite stress in *Bacillus subtilis* as revealed by transcriptomics and redox proteomics. Mol Cell Proteomics. 10(11):M111 009506. doi:10.1074/mcp.M111.009506

Chi BK, Huyen NTT, Loi VV, Gruhlke MCH, Schaffer M, Mäder U, Maass S, Becher D, Bernhardt J, Arbach M, et al. 2019. The disulfide stress response and protein *S*-thioallylation caused by allicin and diallyl polysulfanes in *Bacillus subtilis* as revealed by transcriptomics and proteomics. Antioxidants (Basel). 8(12) doi:10.3390/antiox8120605

Corman VM, Landt O, Kaiser M, Molenkamp R, Meijer A, Chu DK, Bleicker T, Brunink S, Schneider J, Schmidt ML, et al. 2020. Detection of 2019 novel coronavirus (2019-nCoV) by real-time RT-PCR. Euro Surveill. 25(3) doi:10.2807/1560-7917.ES.2020.25.3.2000045

De Maio F, Lo Cascio E, Babini G, Sali M, Della Longa S, Tilocca B, Roncada P, Arcovito A, Sanguinetti M, Scambia G, et al. 2020. Improved binding of SARS-CoV-2 Envelope protein to tight junction-associated PALS1 could play a key role in COVID-19 pathogenesis. Microbes Infect. 22(10):592–597. doi:10.1016/j.micinf.2020.08.006

Deutsch EW, Bandeira N, Sharma V, Perez-Riverol Y, Carver JJ, Kundu DJ, Garcia-Seisdedos D, Jarnuczak AF, Hewapathirana S, Pullman BS, et al. 2020. The ProteomeXchange consortium in 2020: enabling ‘big data’ approaches in proteomics. Nucleic Acids Res. 48(D1):D1145–D1152. doi:10.1093/nar/gkz984

Donma MM, Donma O. 2020. The effects of *Allium sativum* on immunity within the scope of COVID-19 infection. Med Hypotheses. 144:109934. doi:10.1016/j.mehy.2020.109934

Drozdzal S, Rosik J, Lechowicz K, Machaj F, Kotfis K, Ghavami S, Los MJ. 2020. FDA approved drugs with pharmacotherapeutic potential for SARS-CoV-2 (COVID-19) therapy. Drug Resist Updat. 53:100719. doi:10.1016/j.drup.2020.100719

Durfee LA, Lyon N, Seo K, Huibregtse JM. 2010. The ISG15 conjugation system broadly targets newly synthesized proteins: implications for the antiviral function of ISG15. Mol Cell. 38(5):722–732. doi:10.1016/j.molcel.2010.05.002

Fara A, Mitrev Z, Rosalia RA, Assas BM. 2020. Cytokine storm and COVID-19: a chronicle of pro-inflammatory cytokines. Open Biol. 10(9):200160. doi:10.1098/rsob.200160

Gruhlke MC, Hemmis B, Noll U, Wagner R, Luhring H, Slusarenko AJ. 2015. The defense substance allicin from garlic permeabilizes membranes of *Beta vulgaris*, *Rhoeo discolor*, *Chara corallina* and artificial lipid bilayers. Biochim Biophys Acta. 1850(4):602–611. doi:10.1016/j.bbagen.2014.11.020

Gruhlke MC, Nicco C, Batteux F, Slusarenko AJ. 2016. The Effects of Allicin, a Reactive Sulfur Species from Garlic, on a Selection of Mammalian Cell Lines. Antioxidants (Basel). 6(1) doi:10.3390/antiox6010001

Gruhlke MC, Portz D, Stitz M, Anwar A, Schneider T, Jacob C, Schlaich NL, Slusarenko AJ. 2010. Allicin disrupts the cell’s electrochemical potential and induces apoptosis in yeast. Free Radic Biol Med. 49(12):1916–1924. doi:10.1016/j.freeradbiomed.2010.09.019

Gruhlke MCH, Antelmann H, Bernhardt J, Kloubert V, Rink L, Slusarenko AJ. 2019. The human allicin-proteome: *S*-thioallylation of proteins by the garlic defence substance allicin and its biological effects. Free Radic Biol Med. 131:144–153. doi:10.1016/j.freeradbiomed.2018.11.022

Guo NL, Lu DP, Woods GL, Reed E, Zhou GZ, Zhang LB, Waldman RH. 1993. Demonstration of the anti-viral activity of garlic extract against human cytomegalovirus in vitro. Chin Med J (Engl). 106(2):93–96.

Hitl M, Kladar N, Gavaric N, Srdenovic Conic B, Bozin B. 2021. Garlic burn injuriesa systematic review of reported cases. Am J Emerg Med. 44:5–10. doi:10.1016/j.ajem.2021.01.039

Hoffmann M, Mosbauer K, Hofmann-Winkler H, Kaul A, Kleine-Weber H, Kruger N, Gassen NC, Muller MA, Drosten C, Pohlmann S. 2020. Chloroquine does not inhibit infection of human lung cells with SARS-CoV-2. Nature. 585(7826):588–590. doi:10.1038/s41586-020-2575-3

Khubber S, Hashemifesharaki R, Mohammadi M, Gharibzahedi SMT. 2020. Garlic (Allium sativum L.): a potential unique therapeutic food rich in organosulfur and flavonoid compounds to fight with COVID-19. Nutr J. 19(1):124. doi:10.1186/s12937-020-00643-8

Kublik A, Deobald D, Hartwig S, Schiffmann CL, Andrades A, von Bergen M, Sawers RG, Adrian L. 2016. Identification of a multi-protein reductive dehalogenase complex in *Dehalococcoides mccartyi* strain CBDB1 suggests a protein-dependent respiratory electron transport chain obviating quinone involvement. Environ Microbiol. 18(9):3044–3056. doi:10.1111/1462-2920.13200

Loi VV, Huyen NTT, Busche T, Tung QN, Gruhlke MCH, Kalinowski J, Bernhardt J, Slusarenko AJ, Antelmann H. 2019. Staphylococcus aureus responds to allicin by global *S*-thioallylation - role of the Brx/BSH/YpdA pathway and the disulfide reductase MerA to overcome allicin stress. Free Radic Biol Med. 139:55–69. doi:10.1016/j.freeradbiomed.2019.05.018

Masucci MG. 2020. Viral Ubiquitin and Ubiquitin-Like Deconjugases-Swiss Army Knives for Infection. Biomolecules. 10(8) doi:10.3390/biom10081137

Matza D, Badou A, Jha MK, Willinger T, Antov A, Sanjabi S, Kobayashi KS, Marchesi VT, Flavell RA. 2009. Requirement for AHNAK1-mediated calcium signaling during T lymphocyte cytolysis. Proc Natl Acad Sci U S A. 106(24):9785–9790. doi:10.1073/pnas.0902844106

McGauran G, Dorris E, Borza R, Morgan N, Shields DC, Matallanas D, Wilson AG, O’Connell DJ. 2020. Resolving the Interactome of the Human Macrophage Immunometabolism Regulator (MACIR) with Enhanced Membrane Protein Preparation and Affinity Proteomics. Proteomics. 20(19-20):e2000062. doi:10.1002/pmic.202000062

Mehlan H, Schmidt F, Weiss S, Schuler J, Fuchs S, Riedel K, Bernhardt J. 2013. Data visualization in environmental proteomics. Proteomics. 13(18-19):2805–2821. doi:10.1002/pmic.201300167

Minchin WD. 1927. A study in tubercule virus, polymorphism, and the treatment of tuberculosis and lupus with *Oleum allii*. Bailliere, Tindall and Cox: London, UK.110.

Miron T, Listowsky I, Wilchek M. 2010. Reaction mechanisms of allicin and allyl-mixed disulfides with proteins and small thiol molecules. Eur J Med Chem. 45(5):1912–1918. doi:10.1016/j.ejmech.2010.01.031

Miron T, Rabinkov A, Mirelman D, Wilchek M, Weiner L. 2000. The mode of action of allicin: its ready permeability through phospholipid membranes may contribute to its biological activity. Biochim Biophys Acta. 1463(1):20–30.

Müller A, Eller J, Albrecht F, Prochnow P, Kuhlmann K, Bandow JE, Slusarenko AJ, Leichert LI. 2016. Allicin induces thiol stress in bacteria through *S*-allylmercapto modification of protein cysteines. J Biol Chem. 291(22):11477–11490. doi:10.1074/jbc.M115.702308

Münchberg U, Anwar A, Mecklenburg S, Jacob C. 2007. Polysulfides as biologically active ingredients of garlic. Org Biomol Chem. 5(10):1505–1518. doi:10.1039/b703832a

Oguntuyo KY, Stevens CS, Siddiquey MN, Schilke RM, Woolard MD, Zhang H, Acklin JA, Ikegame S, Hung CT, Lim JK, et al. 2020. In plain sight: the role of alpha-1-antitrypsin in COVID-19 pathogenesis and therapeutics. bioRxiv. doi:10.1101/2020.08.14.248880

Ou X, Liu Y, Lei X, Li P, Mi D, Ren L, Guo L, Guo R, Chen T, Hu J, et al. 2020. Characterization of spike glycoprotein of SARS-CoV-2 on virus entry and its immune cross-reactivity with SARS-CoV. Nat Commun. 11(1):1620. doi:10.1038/s41467-020-15562-9

Perez-Riverol Y, Csordas A, Bai J, Bernal-Llinares M, Hewapathirana S, Kundu DJ, Inuganti A, Griss J, Mayer G, Eisenacher M, et al. 2019. The PRIDE database and related tools and resources in 2019: improving support for quantification data. Nucleic Acids Res. 47(D1):D442–D450. doi:10.1093/nar/gky1106

Qin C, Zhou L, Hu Z, Zhang S, Yang S, Tao Y, Xie C, Ma K, Shang K, Wang W, et al. 2020. Dysregulation of Immune Response in Patients With Coronavirus 2019 (COVID-19) in Wuhan, China. Clin Infect Dis. 71(15):762–768. doi:10.1093/cid/ciaa248

Rabinkov A, Miron T, Konstantinovski L, Wilchek M, Mirelman D, Weiner L. 1998. The mode of action of allicin: trapping of radicals and interaction with thiol containing proteins. Biochim Biophys Acta. 1379(2):233–244.

Reiter J, Levina N, van der Linden M, Gruhlke M, Martin C, Slusarenko AJ. 2017. Diallylthiosulfinate (Allicin), a volatile antimicrobial from garlic (Allium sativum), kills human lung pathogenic bacteria including MDR strains as a vapor. Molecules. 22(10) doi:10.3390/molecules22101711

Rivlin RS. 2001. Historical perspective on the use of garlic. J Nutr. 131(3s):951S–954S. doi:10.1093/jn/131.3.951S

Rivlin RS. 2006. Is garlic alternative medicine? J Nutr. 136(3 Suppl):713S–715S. doi:10.1093/jn/136.3.713S

Rossius M, Hochgrafe F, Antelmann H. 2018. Thiol-Redox Proteomics to Study Reversible Protein Thiol Oxidations in Bacteria. Methods Mol Biol. 1841:261–275. doi:10.1007/978-1-4939-8695-8_18

Rouf R, Uddin SJ, Sarker DK, Islam MT, Ali ES, Shilpi JA, Nahar L, Tiralongo E, Sarker SD. 2020. Antiviral potential of garlic (Allium sativum) and its organosulfur compounds: A systematic update of pre-clinical and clinical data. Trends Food Sci Technol. 104:219–234. doi:10.1016/j.tifs.2020.08.006

Schneider WM, Chevillotte MD, Rice CM. 2014. Interferon-stimulated genes: a complex web of host defenses. Annu Rev Immunol. 32:513–545. doi:10.1146/annurev-immunol-032713-120231

Schulte-Schrepping J, Reusch N, Paclik D, Bassler K, Schlickeiser S, Zhang B, Kramer B, Krammer T, Brumhard S, Bonaguro L, et al. 2020. Severe COVID-19 Is Marked by a Dysregulated Myeloid Cell Compartment. Cell. 182(6):1419–1440 e1423. doi:10.1016/j.cell.2020.08.001

Seidel K, Kühnert J, Adrian L. 2018. The complexome of *Dehalococcoides mccartyi* reveals its organohalide respiration-complex is modular. Front Microbiol. 9:1130–1130. doi:10.3389/fmicb.2018.01130

Tian W, Zhang N, Jin R, Feng Y, Wang S, Gao S, Gao R, Wu G, Tian D, Tan W, et al. 2020. Immune suppression in the early stage of COVID-19 disease. Nat Commun. 11(1):5859. doi:10.1038/s41467-020-19706-9

Wang H, Yang P, Liu K, Guo F, Zhang Y, Zhang G, Jiang C. 2008. SARS coronavirus entry into host cells through a novel clathrin- and caveolae-independent endocytic pathway. Cell Res. 18(2):290–301. doi:10.1038/cr.2008.15

Weber ND, Andersen DO, North JA, Murray BK, Lawson LD, Hughes BG. 1992. In vitro virucidal effects of Allium sativum (garlic) extract and compounds. Planta Med. 58(5):417–423. doi:10.1055/s-2006-961504

Wei LL, Wang WJ, Chen DX, Xu B. 2020. Dysregulation of the immune response affects the outcome of critical COVID-19 patients. J Med Virol. 92(11):2768–2776. doi:10.1002/jmv.26181

Wheeler AP, Wells CM, Smith SD, Vega FM, Henderson RB, Tybulewicz VL, Ridley AJ. 2006. Rac1 and Rac2 regulate macrophage morphology but are not essential for migration. J Cell Sci. 119(Pt 13):2749–2757. doi:10.1242/jcs.03024

Zecha J, Lee CY, Bayer FP, Meng C, Grass V, Zerweck J, Schnatbaum K, Michler T, Pichlmair A, Ludwig C, et al. 2020. Data, Reagents, Assays and Merits of Proteomics for SARS-CoV-2 Research and Testing. Mol Cell Proteomics. 19(9):1503–1522. doi:10.1074/mcp.RA120.002164

Zhang L, Lin D, Sun X, Curth U, Drosten C, Sauerhering L, Becker S, Rox K, Hilgenfeld R. 2020. Crystal structure of SARS-CoV-2 main protease provides a basis for design of improved alpha-ketoamide inhibitors. Science. 368(6489):409–412. doi:10.1126/science.abb3405

Zhou P, Yang XL, Wang XG, Hu B, Zhang L, Zhang W, Si HR, Zhu Y, Li B, Huang CL, et al. 2020. A pneumonia outbreak associated with a new coronavirus of probable bat origin. Nature. 579(7798):270–273. doi:10.1038/s41586-020-2012-7

